# Baseline Acute Myeloid Leukemia Prognosis Models using Transcriptomic and Clinical Profiles by Studying the Impacts of Dimensionality Reductions and Gene Signatures on Cox-Proportional Hazard

**DOI:** 10.1101/2022.12.06.519415

**Authors:** Léonard Sauvé, Josée Hébert, Guy Sauvageau, Sébastien Lemieux

## Abstract

Gene marker extraction to evaluate risk in cancer can refine the diagnosis process and lead to adapted therapies and better survival. These survival analyses can be done through computer systems and Machine Learning (ML) algorithms such as the Cox-Proportional-Hazard model from gene expression (GE) RNA-Seq data. However, optimal tuning of CPH from genome-wide GE data is challenging and poorly assessed so far. In this work we propose to interrogate an Acute Myeloid Leukemia (AML) dataset (Leucegene) to derive key components of the CPH driving down its performance and discovering its sensitivity to various factors in hoping to ameliorate the system. In this study, we compare the projection and selection data reduction techniques, mainly the PCA and LSC17 gene signature in combination with the CPH in a ML framework. Results reveals that CPH performs better with a combination of clinical and genetic expression features. We determine that projections performs better than selections without clinical information. We ascertain that CPH is affected by overfitting and that this overfitting is linked to the number and the content of input covariables. We show that PCA links clinical features via ability to learn from the input data directly and generalizes better than LSC17 on Leucegene. We postulate that projection are preferred than selection on harder task such as assessing risk in the intermediate subset of Leucegene. We extrapolate that these findings apply in the more general context of risk detection via machine learning in cancer. We see that higher capacity models such as CPH-DNNs systems can be improved via survival-derived projections and combat overfitting through heavy regularization.

**Author summary:** This study aims to investigate the feasibility of using gene expression to evaluate risk in cancer, and to compare the projection and selection data reduction techniques. The study used the Leucegene dataset to compare the PCA method and a previously published 17 genes signature in combination with the Cox-Proportional-Hazard model in a machine learning framework. Results showed that CPH was affected by overfitting and that this overfitting was linked to the number and the content of input covariables. The study found that PCA links clinical features via ability to learn from the input data directly and generalizes better than LSC17 on Leucegene. The study concluded that projections are preferred than selection on harder task such as assessing risk in the intermediate subset of Leucegene and can be tuned to improve their performance.

**Data availability statement:** Source code for pipelines and algorithms, as well as gene expression matrices, are available here: https://github.com/lemieux-lab/dimensions_coxph. Access to the Leucegene cohort’s survival times can be granted upon request and following ethical review.

## Introduction

Acute myeloid leukemia (AML) is a lethal disease among adults with chances of survival estimated around 28%, 5 years after diagnosis. In the past years, research has allowed the discovery of multiple genomic markers such as the NPM1, FLT3-ITD, IDH1 and other mutations that indicate good or poor prognosis of the disease. Stratification systems based on genomic abnormalities and mutational profile exist through the European LeukemiaNet (ELN) classification (Döhner et al., 2017) and others (Papaemmanuil et al., 2016). So far, these classification strategies allow to divide patients into adverse, intermediate and favorable risk categories. These assessments influence the therapeutic strategies that a clinician would use to treat AML, mainly through chemotherapy alone or combined with bone marrow transplants. Gene expression prognosis markers for AML have also been identified such as HMGA2 expression (Marquis et al., 2018) or the 17-genes stem-cell signature (Ng et al., 2016). The potential of genetic expression and genomic profiling data as a tool for AML prognosis has been well established so far, but the multiplicity of these markers seems to introduce confusion on how to best combine them in order to achieve an optimal stratification. The less prevalent mutations or rearrangements that are now being identified are difficult to exploit for prognosis since cohorts would tend to present few cases for each. Gene expression profiles provide a characterization that is closer to the phenotype could provide a de facto integration of the mutation landscape of a patient sample that would be more convenient for deriving survival models of the disease.

For this reason, initiatives like the Cancer Genome Atlas AML (TCGA, n = 672 samples) and Leucegene (n = 452 samples) have put together AML whole gene expression profiling datasets with clinical annotations to support the creation of such computational approaches. A common survival model is the Cox-Proportional-Hazard (CPH) model (Cox, 1972) of relatively simple design and its ability to manage censored data (Kleinbaum & Klein, 2012a). Unfortunately, gene expression profiles provide a very high dimensionality representation of patient samples that is induced by the quantification of over 20,000 genes. The performance of CPH is negatively affected by an increase in the dimensionality of the input. In the machine learning (ML) literature, this effect is notoriously known as the “curse of dimensionality”. Typically, the more features are made available to a model, the more samples are required to fill the input space. Models also tend to be more prone to overfit as the models’ number of parameters increases with the dimensionality of the input space and rapidly outmatches the number of training samples available (Bengio, Yoshua, Ian Goodfellow, and Aaron Courville, 2017). To reach good performances when using gene expression (GE) profiles, a dimensionality reduction step is typically introduced before optimising a CPH model, thus reducing the negative impact due to the curse of dimensionality. However, factors like the input information density, size of dataset, type of dimensionality reduction technique used and their impact on the accuracy of the CPH is poorly understood. Two distinct data reduction strategies seem to be promising. The gene selection strategy, as used for LSC17 signature based on *stem-cellness*, is proven to predict survival well in some test cohorts. However, an issue that arises when using a gene signature as the basis for a survival risk score is the sheer number of possible signatures. A well-documented study clearly displaying this issue can be found in (Boutros et al., 2009) where the authors evaluated all 6-gene signatures and found that an important number of non-overlapping signature had high level of performance. Here, we will explore larger signatures through random sampling and assess their performance with respect to a carefully assembled signature. As an alternative to this gene selection processes, we hypothesized that informed projection algorithms such as the principal component analysis (PCA) should increase the performance of survival models and provide a more systematic way to build survival models from GE data.

In this work, we want to test the benefits of using data-driven projections and test if CPH models can be trained to predict risk in AML. To do so, we will use the Leucegene dataset to test different combinations of input representations (either selections or projections) with CPH. We want to frame this work in a ML setup, meaning that performances will be assessed on unseen data through internal cross-validation. In parallel, we want to confirm that the predictive signal in gene expression profiles is highly distributed across the transcriptome by assessing the performance of random gene signatures as compared to biologically informed LSC17 signature. Finally, we will confirm our findings through the analysis of an external dataset (TCGA) and an *a priori* more challenging subset of the Leucegene cohort.

## Methods

### Datasets

#### Leucegene

452 public AML samples Hi-Seq Illumina RNA-seq data. Geo accession: GSE67040. Raw Fastq alignment and read counts were computed with Star-RSEM. Read counts are reported as transcripts per million (TPM) normalised and log_10_(*x* + 1) transformed. The reference genome version is GRCh38. Clinical annotations are obtained through the Banque de Cellules Leucémiques du Québec (BCLQ). We chose to restrict the size of the GE profiles to 19,183 expressed protein coding genes, plus 17 important clinical features (i.e. sex, NPM1, FLT3-ITD, IDH1 mutations, age at diagnosis, adverse, intermediate or favorable cytogenetic risk groups and the WHO classification). The set of genes for the random projections and random selections was further reduced to the 8,591 genes (50%) with highest variance. From this dataset, two subsets were derived: 1) Prognostic subset (n = 300, censored = 77 (~26%)): This subset only retain samples from patients that underwent intensive therapies. Maximum follow up is 6,000 days (16 years) after diagnosis. 2) Intermediate subset (n = 177, censored = 40 (~22.5%)): From the prognostic subset, only patients classified as intermediate cytogenetic risk were considered.

#### TCGA

Data from the Cancer Genome Atlas Acute Myeloid Leukemia projects (BEAT-AML, TARGET-AML) was obtained, representing a total of 672 samples (Tyner et al., 2018). Only 140 samples were identified that had both RNA-seq data and complete survival information. Consequently, this cohort is very similar to the one used by (Ng et al., 2016) to validate the LSC17 signature. Gene counts and clinical annotations were downloaded from raw files via the *cBioPortal for cancer genomics*. In this subset of TCGA (n = 140, censored = 55 (~39.2%)), the maximum follow up is 2,200 days (6 years) after diagnosis.

### Dimensionality reduction

To perform and test the different dimensionality reduction techniques we selected two projection-based (random and PCA) and two selection-based algorithms (random and LSC17).

#### Random projection

The random projection we use is the random projection using gaussian distribution as described and explored in (Fradkin & Madigan, 2003). We feed the random projection algorithm a filtered gene list for which expression variance is higher than the median of variance of all genes (50%). This helps Cox-PH optimize better, by removing low varying genes or unexpressed genes that the CPH handles poorly. The random projection algorithm implementation is done via the python *sklearn* random projection method. To get a robust metric of the performance of this technique, we sampled 1,000 projections with increasing size (1-50 dimensions).

#### Principal Components Analysis (PCA)

As a data-driven projection, we use the PCA as it performs a linear transformation of the input data by rotation. Unlike the random projection, the PCA attempts to disentangle principal factors in the input by returning a list of eigenvectors ranked according to the variance they explain in the input. By projecting the original data on the eigenvectors, we get a compressed view of the data that preserves its general structure. A more complete description of the internal computations of the approach can be found at section 5.1.9 of (Bengio, Yoshua, Ian Goodfellow, and Aaron Courville, 2017). The implementation of the PCA algorithm was conducted using python *sklearn linear_decompostion* PCA method with default parameters.

#### Random selection

We use a simple random selection-based dimensionality reduction by selecting genes randomly without replacement. By sampling 1,000 gene signatures at each tested input dimension (1-50), we get the distribution of performance for this class of dimensionality reduction.

#### Informed selection

As a biology-derived signature or informed selection-based method, we choose to compare the dimensionality reduction policies presented before with the LSC17 (Ng et al., 2016), a 17-genes signature demonstrated to be of good prognostic value in AML. The LSC17 gene set is composed of the following genes: CD34, LAPTM4B, ARGHGAP22, LOC284422, KIAA0125, MMRN1, DNMT3B, SOCS2, CDK6, NGFRAP1, CPXM1, ZBTB46, DPYSL3, NYRIN, GPR56, AKR1C3, EMP1. By suspecting that stem-cell-like cells display resistance features to chemotherapy in AML, authors of this paper constructed a microarray dataset from 78 patients using xenotransplantation to compare 138 LSC+ and 89 LSC-samples that were used to derive a list of genes linked to stemcell patterns. This list was then manually and computationally enriched and filtered for genes that have prognostic value in various datasets such as TCGA.

### Clinical feature correlations and prediction

To quantify the relationship between gene expression and clinical features, we evaluated the strength of the correlations with two metrics: Pearson moment correlation (*numpy’s corrcoef* method) and logistic regression performance. A classification task was set in place to train a logistic regression from gene expressions to predict clinical features. The parameters were optimised using the *sklearn* implementation with L2 regularisation parameter C set to 0.99, and default hyper-parameters. The area under the Receiver-Operator-Curve (AUC) from *sklearn* is then used as a performance metric.

### Survival analysis and evaluation

#### Cox-Proportional Hazard (CPH)

For this work, we used the CPH survival model (Cox, 1972) to test the different data projections described above for the context of survival prediction. We are aware that a large variety of more recent models exist based on combination of CPH and deep neural networks (CPH-DNN), they were introduced in the mid 90’s (Faraggi & Simon, 1995) and further refined more recently in implementation such as DeepSurv (Katzman et al., 2018) and Deep-Hit (Lee et al., 2020). We used the python package *lifelines* Cox proportional hazard method (Davidson-Pilon, 2019) with the L2 regularisation parameter set to 10^-10^, a very small amount as it seemed not to affect performance drastically. Note that the CPH is an interpretable model because we can relate the input factors directly and interrogate the strength of the correlations with statistical tests. In this work, we will not explore this feature. Instead, we use the CPH as a benchmark survival model to test the impact of different projection and selection methods.

#### Kaplan-Meier (KM) plots and log-rank test

To compare the performances of the trained models, we stratified our cohorts into three risk group categories (intermediate, favorable, adverse) following the original distribution of risks in the Leucegene and TCGA cohorts respectively. The assignations of risks were done using the CPH risk scores obtained through cross-validation. The KM (Kaplan & Meier, 1958) plots provide visual assessment of the resulting risk groups. Log-rank test is applied to the data presented in figure 4b,c to compare the survival of the two resulting groups in the Leucegene intermediate subset (Kleinbaum & Klein, 2012b). A p value < 0.01 allows to dismiss the null hypothesis that groups have same survival. We used implementations provided by the *lifelines* package.

#### Concordance index

We used the concordance index to evaluate our CPH models. The advantage of this method is that it evaluates the model performance without the need to set a threshold beforehand. In this regard it behaves similarly to the AUC in classification. The concordance index is the probability that each sample is ranked properly according to the other samples, it spans values between 0.5 for random and 1 for perfect ranking, taking into consideration censored values. Preliminary analyses reflected that this metric fluctuates as the number of samples decreases in the context of crossvalidations.

#### Cross-validation and bootstrapping

Two cross-validation strategies were used. In the first setup, we proceeded by 10-fold crossvalidation. Then, out-of-training risk scores for the whole cohort were used to compute a c index. 95% confidence interval were computed using 10,000 permutations with redraws of the risk scores. In the second setup, 1,000 random projections and 1000 random selections were computed with increasing sizes (from 1 to 50) and evaluated by 5-fold cross-validation, distributions are reported as histograms. For the PCA with increasing input dimensionality and LSC17, 1,000 random shuffles followed by 5-fold cross-validation procedure were applied to obtain distributions of equal length to the random. Note a bias exists against c index reported through the 5-fold crossvalidations as these models are trained on 80% of the samples instead of 90% in the first setup.

## Results

### Evaluation of Cox-PH with different input types

In order to compare the effect of different input types for a Cox-PH and to determine which strategy is the best for training a survival model from gene expression profiles, we put in place the benchmark displayed in figure 1. This also provides useful insight on the repartition of risk groups within the Leucegene cohort. The first model we present is a visual benchmark based on cytogenetic risk as the pre-existing model. Then, the reference CPH on clinical features model is the CPH base reference.

**Figure 1:**
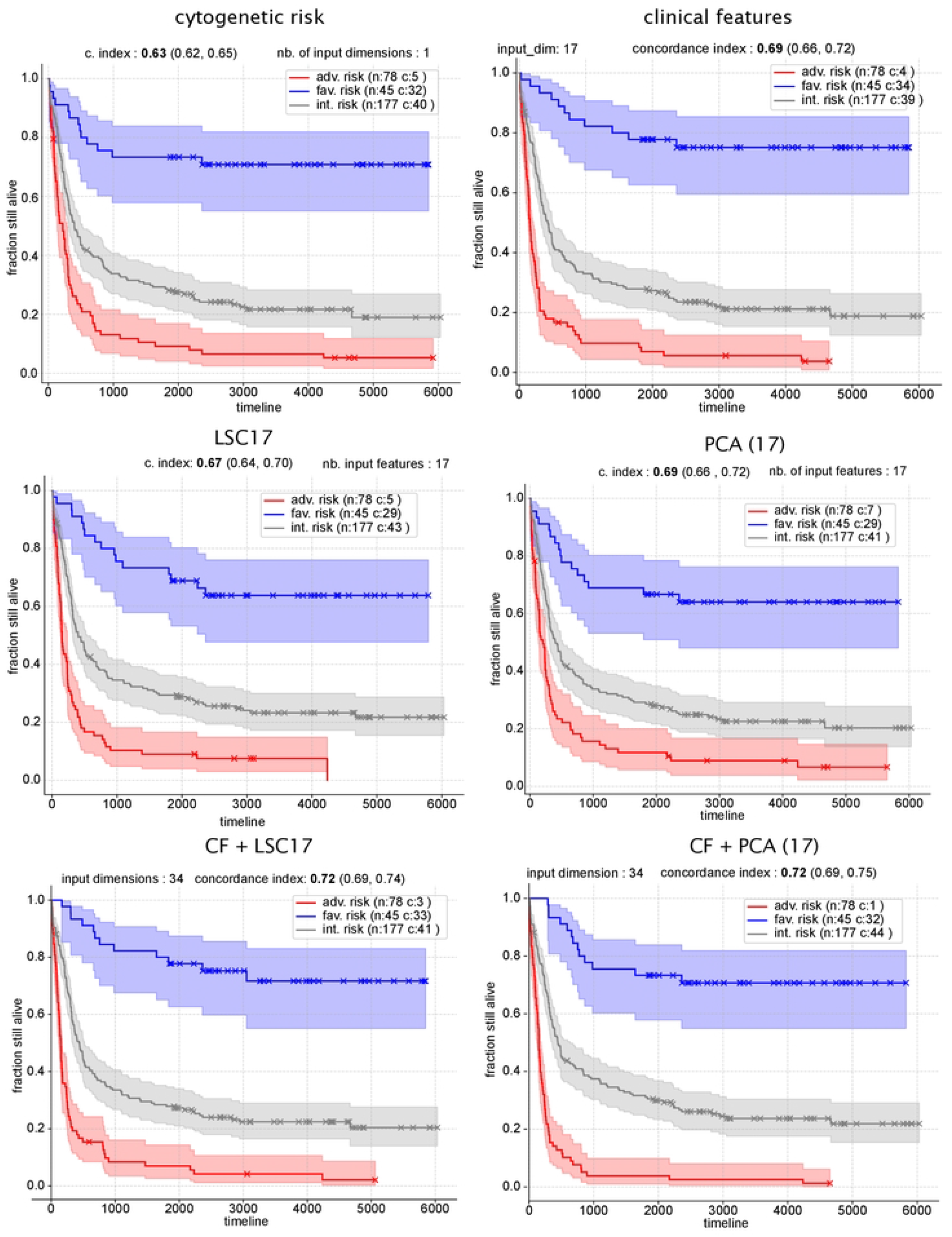
Predicted survival curves in Leucegene dataset with Cox-PH models with clinical, LSC17 and PCA input features. CPH testing performance is obtained using 10-fold cross-validation concordance index and bootstraping for 95% (min, max) confidence interval. y-Axis: survival / fraction still alive. All models presented here are Cox-PH regressions with 12 regularization trained and tested internally on the Leucegene cohort (n: 300, censored: 77) with lifelines, and varying input features, **cytogenetic risk**: repartition according to cytogenetic risk, **clinical features**: features relative to the WHO classification (11 AML subgroups), **cytogenetic risk**: favorable, intermediate or adverse, NPM1, FLT3-ITD & IDH1-R32 mutations, sex and age. total: 17 dim. input. **LSC17**: gene signature expression, 17 dim. input. **PCA (17)**: Gene expression data projected onto its PCA’s first 17 components. **CF-LSC17**: clinical and LSC17 feratures. 34 dim. input. **CF + PCA (17)**: clinical features and gene expression features through PCA. 34 dim. input. All curves are fitted with *lifelines* Kaplan-Meier survival curves.

#### Overall accuracy of systems

All the tested inputs model survival better than using cytogenetic risk only. Models tend to have moderate accuracies by c index, never reaching 0.75. Although we cannot exclude that this performance is an upper bound, many strategies are yet to be attempted to improve performance and performance could still be limited by the limited size of cohorts. In this section, we investigate the impact of the input type (GE or CF) on performance.

#### Model accuracy comparisons

Among all tested models, best performance is achieved with the multivariate models using the 17 clinical features and 17 gene expression components through PCA or LSC17 both displaying concordance indices of 0.72. Looking at models that were trained only on clinical features (CF) or gene expression features (LSC17 or PCA(17)), models using a CPH on CF or PCA yield the best performance by cross-validation using the Leucegene with a concordance index of 0.69. Highest performance with clinical factors is expected since the cytogenetic risk explains most of the survival in the cohort as shown in the upper left panel of figure 1. However, the fact that PCA(17) captures as much information as the clinical factors model is slightly surprising and hints that expressions are as good prognosis markers as clinical factors are and could be used for prognosis. PCA(17) slightly outperforms LSC17 for determining risk in this setup by concordance index (0.69 vs 0.67). PCA(17) captures well clinical features such as the cytogenetic risk which itself correlates the most with survival in the cohort. To put it all together, we propose that PCA and clinical features share great similarities and that LSC17 determines survival, but with slightly less precision than the PCA. Furthermore, we pose that LSC17 and CF share less similarities than PCA and CF. Consequently, the fact that LSC17 + CF determines survival with the best accuracy might be explained by LSC17 carrying information complementary to clinical features.

### Correlations between gene expressions and clinical features

In this section, we wanted to verify how well principal components (PCs) align with clinical features. To do so, we computed the Pearson-moment correlation between the first 30 principal components of the PCA and the clinical features directly (Fig. 2A). By design, PCs are ranked according to explained variance in the original GE matrix. We observe variable correlation strengths among first PCs and clinical features. The first 10 PCs tend to correlate more to the cytogenetic risk than later components. The NPM1 and FLT3-ITD mutational status correlate strongly with several of the first 6 PCs and sex appears to be capture by later PCs (13-22). However, it seems that some PCs do not correlate strongly with any feature, and it is puzzling to see this for the first two PCs. It might be possible that these components capture features of the patient samples are not present in the clinical features.

**Figure 2:**
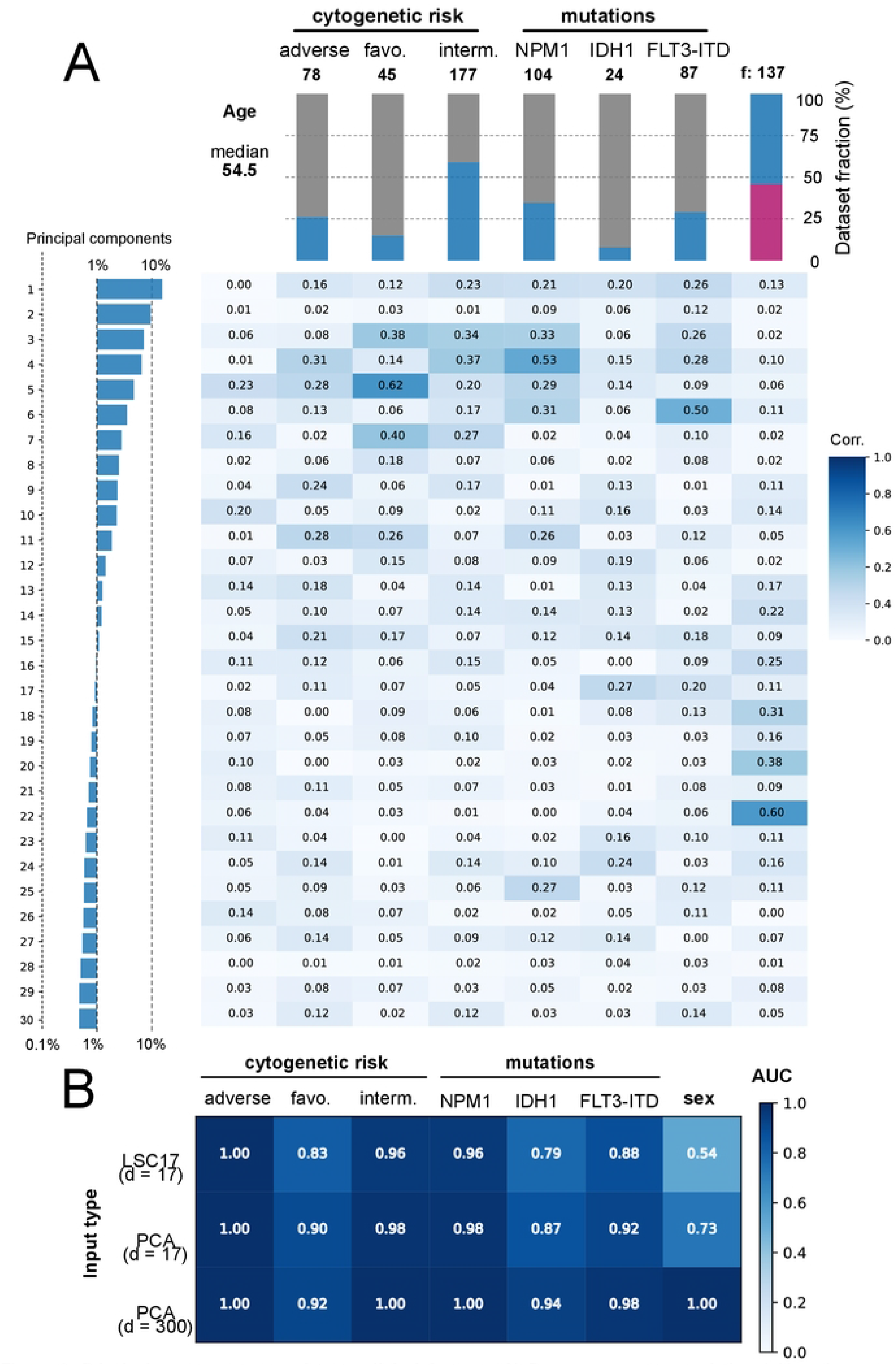
Principal components correlate to clinical features. **A) Pearson-moment correlation** (with python *numpy* package), heatmap of the absolute value as the sign of corn is irrelevant here. Y axis: principal components, X axis: clinical features. Left panel: explained variance in on log scale for clearer discrepancies. Dashed line are 1% variance. 10% variance. Top panel: incidence of target features, by absolute number. Bars: fraction of dataset (n=300). **B) Prediction performance** (ROC-AUC) of logistic regression classifiers (*sci-kit learn*) by feature from LSC17 gene expression input, PCA (d = 17) with 17 first components and PCA (d = 300) with all PCs by leave-one-out crossvalidation.

### LSC17 and PCA provide strong predictors of clinical features

Although individual PCs correlate at diverse degree to clinical features directly, we wanted to ascertain the capacity of multiple PCs to determine single clinical features together. For this, we used a logistic regression with regularization to predict each clinical feature (Fig. 2B). We observe that with equal number of input features, PCA performs better than LSC17 at every task. This confirms that PCA captures the CF better than LSC17. The fact that PCA(17) better predicts sex than LSC17 is simply a consequence that the PCA aims to capture variance within the dataset without a priori knowledge of the underlying biology. PCA with all 300 PCs determines cytogenetic features very well and determines mutations and sex better than the other methods.

### Impact of input space dimensionality on Cox-PH performance

Intuitively, adding more input data to the model should increase its performance, until the increased number of free parameters in the model leads to an overfit of the training data. To quantify these effects, we varied the number of input features extracted from the PCA (Fig. 3). In parallel, we wanted to test the benefit of using data-driven data reduction algorithms as compared to either random projections or random selection with an identical number of free parameters. Our goal is to deconvolute the impact of the data-driven process during the data-reduction step. We used LSC17 as a static performance baseline since we cannot increase nor decrease the size of this pre-determined signature. The effect of an increasing number of PCs is shown both with and without CF.

**Figure 3:**
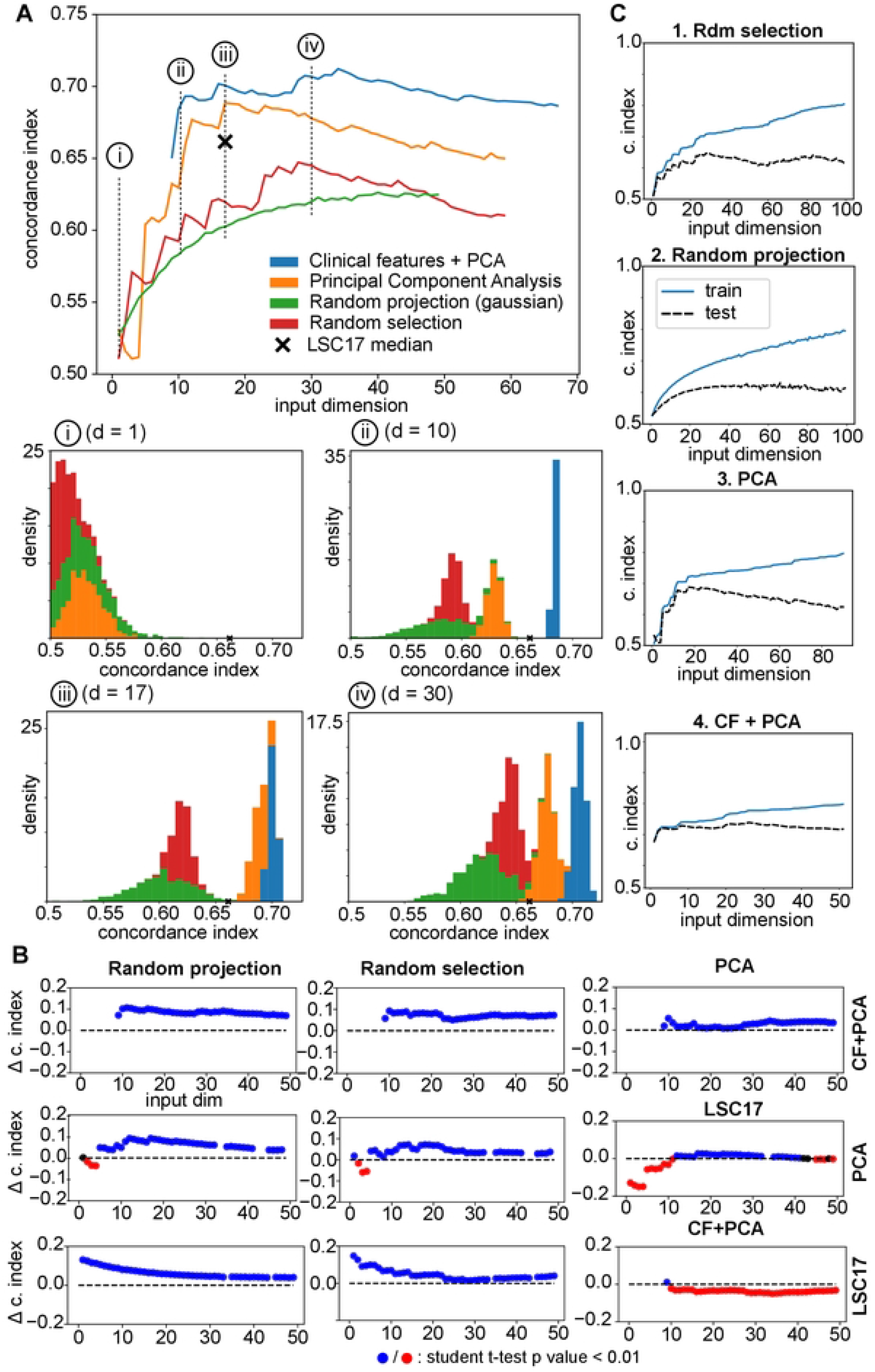
Analysis of Cox-PH with increasing number of input clinical and gene expression features. **A) dimensionality impacts the Cox-Proportional Hazards model**. Performance of Cox-PH by 5-fold cross-validation by number of input features. Panels i, ii, iii, iv display the bootstraped concordance index distributions by projection type for input dim = 1, 10. 17, 30. **B) Performance differences of projection and selection types by dimension** displayed by c. index distribution mean differences and Student t-test p values by input dimension between projection types. Computed differences is mean of distribution projection method 1 (horizontal) minus mean distribution of projection method 2 (vertical). **C) Cox-PH is sensitive to overfitting**. Overfitting curves: training and testing performance for different projection or selection types. Beyond certain dimensionality, performance on training still increases and performance on test decreases.

We first observed slight, but systematic, performance loss from switching from a 10-fold crossvalidation scheme to a 5-fold cross-validation (trained on 80% of the samples). This does not affect our capacity to compare the various input representation within the same cross-validation scheme. PCA, CF-PCA and random selection display the same performance trends, namely increasing by the number if input dimension until a peak is reached, then undergoing a constant decrease in performance as the number of input dimension becomes higher (Fig. 3A). The random projection algorithm displays a slightly different trend, showing a constant but minimal increase in performance by the number of input dimensions and then stabilizing towards a c index of 0.6 on average at about 30 dimensions. CPH on CF-PCA outperforms all other methods at any dimension (Fig. 3B). Early PCs (1 to 5) appear to encode information irrelevant to prognosis and, as such, display less performance than LSC17. PCA at 10+ dimensions have better performance than LSC17 and random selection until high dimensionality is reached when performance decreases. CF-PCA vastly outmatches PCA at low dimensions, but after 17+ dimensions, difference becomes very thin. LSC17 outmatches random selection at low dimensions, but as dimensionality increases this margin also gets very thin. Random selections at high input dimensionality almost get same performance as a data and biology-driven selections such as LSC17. We observe that the difference of performance between the two strategies gets very small near 0 at around random selections of 29 gene expressions (Fig. 3B, lower middle panel). The surprisingly high performance of larger random signatures (e.g., 29 genes) is likely due to a high probability to include genes that correlate either directly with clinical features, genes of the LSC17 or PCs. However, at equal number of dimensions, random selections of 17 genes never surpass LSC17 performance. The best approach seems to combine clinical features and highly informative components of data-driven projections such as a PCA. Random selections perform better than random projections in general. All methods display features that indicate overfitting as the number of input dimensions is increased (Fig. 3C). Random projections, selections and PCA all share the advantage that their number of input dimensions can be tuned at will while fixed length signatures such as LSC17 are more difficult to adjust when new cohorts become available.

### Overfitting studies

Overfitting is a common theme when training machine learning models with very high number of free parameters on small cohorts, it occurs when the algorithm memorizes the training data rather than extracting general features. This phenomenon can be directly observed on models as an increase with capacity of the difference between training and test performance. Figure 3C shows both performance measures on the model trained for figure 3A.

All methods appear to suffer from overfitting. As suspected, for every method as input dimension increases overfitting increases, meaning that the difference between the training and testing performance increases. However, the exact number of dimensions after which models start to overfit varies. For the random selection, random projection and PCA policies, difference between testing and training performance is notable as early as the first two or three dimensions. However, testing performance still increases at this point and is not considered as being in the overfitting regime. For the random selection scheme, the point of decrease happens around 27 dimensions. Even though training performance continues to grow as we add more and more dimensions, testing performance starts to drop slowly. This overfitting point happens around dimension 17 for the PCA model and the overfitting point seems to be absent with the random projection scheme, which seems to present slightly different trends regarding testing performance. Indeed, this scheme seems to provide steady growth in testing performance as we add more input dimensions, but the performance is always at the lowest levels. The difference between training and testing set performance is also markedly high at higher dimension. It raises the question whether the performance of the random projection should be the lower plateau for the performance of the Cox-PH in this context. The CF+PCA suffers less overfitting. The overfitting threshold seems to be located much farther than other methods, standing around 35 input dimensions. After this point testing performance starts decreasing as do the other methods. We postulate that the fact that the model uses clinical features delays the overfitting phenomenon in this experiment. It is supported by the fact that PCA alone suffers more overfitting than when added to CF. As later PCs get more and more noisy, CPH struggles to deconvolute that noise from general properties of the data necessary to be optimized for risk detection. CPH suffers from overfitting regardless of the nature of its input and the burden of overfitting variates according to the content of the input data itself with Cox-PH. Providing insightful input may delay the loss of generalization power. It might also be influenced by the size of the sampling.

### Validations on external data and unclassified subset of Leucegene

Previously, we determined that PCA with cox-PH predict survival at a good level on the whole cohort. Here, we wanted to test if the PCA method can decipher risk harder contexts i.e., with a cohort with uncertain clinical annotations such as the intermediate category. To compare, we use a selection-based method of data reduction LSC17 prognosis gene signature. The reason we use the intermediate cytogenetic risk subset is that we think there may be factors hiding inside this cohort that may allow us to pose higher accuracy diagnosis for newly acquired samples of unassigned risk category or assigned with uncertainty. We also wanted to apply this method to an external database, data from the Cancer Genome Atlas (TCGA) Acute Myeloid Leukemia samples. This dataset had to be filtered because the survival information was missing or invalid and after processing this data contains about 140 samples. Like the Leucegene dataset, many samples are right-censored (39%). Because of its smaller size and its censorship rate, we consider that this data is noisier than our original dataset.

Because we impose our models some new constraints (lower n, higher noise, uncertainty in cohort) we expect a lower overall performance than to the original setup, which is verified with the results collected. Note that setups differ from experiments displayed on figure 4a, d vs 4b, c, e, f and cannot allow direct performance comparison. From results on figure 4 we also conclude that gene expression determines survival in difficult context, when clinical information is missing, and data is noisy because both methods can separate the high-risk and low risk samples efficiently (p value of log-rank test < 0.01).

**Figure 4:**
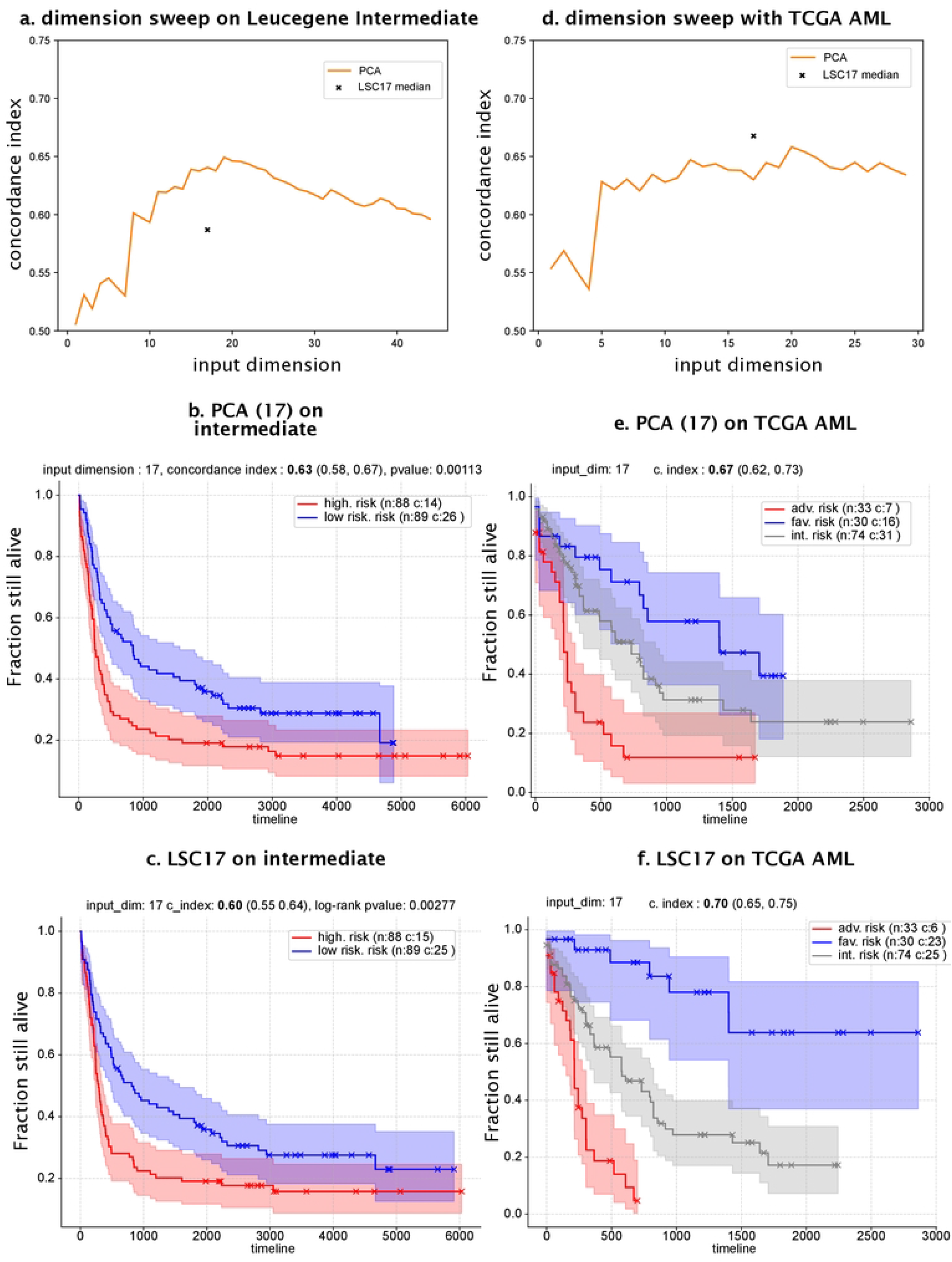
Predicted survival curves on from gene expression with Cox-PH models using PCA and LSC17 gene signature on intermediate subset and TCGA without clinical information. Multivariate Cox-PH evaluation using 5-fold cross-validation c index by bootstraping for 95% (min, max) confidence interval. y-Axis: predicted survival ? fraction still alive, **(a,b,c)** Leucegene intermediate cytogenetic risk cohort (n=177) separated in lowest 50% and 50% highest risk based on predicted scores, **(d,e,f)** AML expression and survival data from the Cancer Genome Atlas (TCGA). Cox-PH regression models were trained with *lifelines* on Principal component analysis decomposition of gene expression features and with LSC17 gene selection methods. Kaplan-Meier survival curves and logrank tests computed with *lifelines*.

While PCA performs better than LSC17 (concordance index = 0.63 vs 0.60) on Leucegene intermediate subset, PCA is outperformed by LSC17 on TCGA (0.70 vs 0.67). This behavior is hard to explain, and we cannot conclude with certainty which strategy is the best to use in this precise context. TCGA is a smaller set of samples with higher censored survival times. LSC17 was developed and validated with the TCGA data. We also think it is possible to tune the number of dimensions used by the CPH from PCA and increase the performance of the method (as shown on figure 3.d). With the Leucegene intermediate samples, PCA succeeds in retrieving signal from the gene expression related to survival as it is unclear whether the LSC17 signature has captured any information or simply noise. This is explainable that the fact that PCA had the possibility to learn from the intermediate cohort, as LSC17 was pre-trained on another set of samples, thus generalizing badly on this harder subset. Surprisingly, high-risk low-risk separation is significative according to the p value of the log-rank-test with LSC17 on the intermediate. This discrepancy between the accuracy through c index or Kaplan-Meier fitting and log-rank-test is hard to explain in this setup.

## Discussion

### Random vs informed

LSC17 as one of many random selection LSC17 is good but can be surpassed by random in some cases. One of the major points of this study is to verify the fact that transcriptome profiling data exhibit highly repetitive information. We verified this by highlighting minor differences in performance of large random signatures and thinner more refined and informed gene selections such as LSC17. Although many genetic components linked to survival could coexist in the data, we rather support the opposing idea that very few components explain survival and that very many genes (all?) are correlated to those very few components. With that idea we suspect that LSC17 captures an optimal number of genes that represent a large portion of the survival predictivity of the gene expression matrix. However, the high codependence of genetic information poses a challenge with machine learning processes and must be dealt with. We propose that instead of being restricted by this feature of gene expression data we rather utilize this fact to our advantage to increase performance. For this task, data-driven projections that derive the main components and perform data reduction simultaneously present promising venues. While random projections use this idea, they seem to perform badly with Cox-PH. PCA on the other hand seems to take profit of this idea and can get good performance at survival prediction with Leucegene and TCGA. Adding informative input or informed projections to the Cox-PH also makes it less sensitive to the increasing size of the input, which makes the Cox-PH more robust to overfitting. Random processes are insufficient in this case in themselves to explain as much information as an informed projection. LSC17 is an informed selection by design and can be considered a good signature compared to the average of randomly selected genes. But at a size large enough to correlate to all the key genetic components that explains survival well by chance, some rare random signatures perform better than LCS17. This reinforce the idea that even powerful signatures are repetitive along the transcriptome brought by (Boutros et al., 2009) and inspired this methodology.

### Projection vs selection

One of the topics of our research aimed to determine the optimal dimensionality reduction strategy of whole transcriptome profiling data for survival analysis with Cox-PH. This aimed to determine the best policy between selection-based and projection-based reduction techniques. Results presented in the previous sections helped us shed some light on this question by highlighting key aspects of both strategies regarding performance in survival modeling. Data-driven projections such as PCA tends to capture clinical features better than selections. PCA has the major advantage of being tunable by the number of input PCs fed to the model. Random selection processes also have this feature but tend to be outperformed easily by data-driven projections at same dimensionality. PCA captures information that the random projection does not and that hinders the CPH model. The tested data-driven selection LSC17 performs well on our benchmarks compared to random selections but have poor performance compared to the projection method PCA and the multivariate model (clinical features + genetic features). On harder tasks i.e., when cytogenetic risk information is missing (Leucegene intermediate cohort) or when the number of samples is reduced and censorship is high, PCA seems to perform well. The LSC17 was largely outperformed by PCA on the intermediate subset, sign that PCA is a more generalizable approach. The fact that LSC17 performs as well as PCA on TCGA indicates that selection-based approaches are still powerful in some cases, but the conditions necessary for their high performance are hard to extract with our current data. We think that data-driven projections such as PCA should be preferred when building GE survival analysis models. These models can be tuned by the optimal input dimensions and tend to be more generalizable than random or data-driven selections-based signatures. Perhaps Leukemia SC (stem-*cellness*) plays less a central role in Leucegene dataset and PCA captures survival better in that case.

### Improvements to the systems

The results presented in this work have shown that elementary linear computer systems (CPH+PCA) can model survival well. So far determining the upper bound on the performance on survival prediction using Leucegene is hard. A point can be made that perhaps the upper bound of performance has already been reached with the empirical results shown so far, and that closing the margin between the current best concordance (0.71) and perfect performance (1) on Leucegene is impossible due to the unmanageable noise inherent to survival data. Survival data contains censored data (missing values), and gene expression data is massive and hard to handle. Although this is possible, we argue that it is unlikely the case. Increasing the upper bound of the performance of survival models from GE data should be possible by addressing the limitations of the current models. Limitations inherent to CPH concern the curse of dimensionality linked directly to the size of the cohorts and the dimensionality and the informativeness of the input and overfitting. Also, because the CPH is linear, it should be limited by capacity if non-linear interactions are indeed present in the data. It is also unclear if the Newton-Raphson optimization of CPH through Cox-Negative-Likelihood loss that indirectly correlates the evaluated concordance index is the correct approach for optimal performance. We also identified limitations to the projection approaches fed to the CPH such as the PCA, as it may contain components in its first PCs that do not correlate to survival. Also, later PCs contain low explained variance components. Many improvements to the current systems can be made to obtain risk evaluation systems that perform better than the models presented here.

Increasing the number of samples for training the system should affect positively the performance. We can observe a loss of performance between the PCA model on Leucegene full vs the smaller intermediate cohort. Raising the number of training samples should have the opposite effect on performance. Determining the optimal number of samples in the training cohorts is currently impossible to ascertain. Methodologies able to estimate this number would help survival ML from GE data training tremendously.

From past empirical and theoretical results increasing the capacity of the CPH model should improve its accuracy. A CPH-deep-neural network architecture was first introduced by (Faraggi & Simon, 1995) and was implemented with DeepSurv (Katzman et al., 2018). Deep-neural network models model non-linear interactions. Modern computational resources manage the training of these models. As (Katzman et al., 2018) demonstrated, these models capture survival from non-linear input space better than the original linear CPH on simulated data. Transforming the linear CPH to a DNN has the downside of removing the interpretability of the system i.e., our capacity to relate the model’s parameter back to the input directly. Although, in the quest for modeling with the best performance, this loss of interpretability is irrelevant. However, increasing capacity freely might not automatically affect positively performance. From the results treated here regarding generalization power all CPH models suffer heavily from overfitting even with nondeep architectures. In this line of thought, higher capacity models such as CPH-DNN should overfit even more.

Another idea derived from the principles discussed in this work is to drive an internal projection method directly to optimise survival. In that way, data representation and dimensionality reduction are done with the objective of modeling survival. Models presented so far compartmentalised the data reduction step (PCA) and the survival function optimization step (CPH tuning). Plus, PCA performs linear transformations, which can be possibly insufficient to model data. Neural networks such as t-SNE (Hinton & Van Der Maaten, 2008) and Factorized Embedding (Trofimov et al., 2020) can learn useful non-linear data-driven representations. Adaptation of these DNNs in practice to learn survival-directed representations with a CPH could theoretically improve performance.

Some features of the random selection process might be beneficial to a CPH. The highly repetitive nature of the gene expression data allows random projections at high dimensionality to capture almost the same amount of information as directed thinner selections (LSC17). This hints to the fact that models could be able to learn where the important features are in the input through attention networks mechanisms. These computational structures were originally inspired from human visual attention mechanisms (Itti & Koch, 2001). Networks that implement those ideas mostly have been applied to image classification (Zhao et al., 2017) (Wang et al., 2017) via convolutional layers in very deep structure. Transposing these ideas to gene expression data is theoretically possible, but whether the structure works in practice is yet to be determined. Attention network mechanisms would gain the benefit of random selections while making them more selective and informed by selecting individual components in the input without transforming them per se and could improve accuracy.

Developing robust regularisation for low and high-capacity CPH models is paramount. So far, regularization has been limited to manual strategies by keeping a low number of input dimensions or by tuning the weight decay hyper-parameter. Note that that these parameters influence each other, as regularization of the system allows input of greater dimension. It would be interesting and probably beneficial to introduce automated regularisation mechanisms. For genomic analysis with “fat” data, whose number of input variables largely exceeds the number of training samples, diet-networks structures have been proposed (Romero et al., 2017). In practice, such a structure would train a network to regularise the parameters of the model and training the objective function simultaneously. Whether such structures can be implemented with transcriptomic profiling data with survival modeling remains to be tested, but the similarity between fat genomic data and transcriptomic data hints that positive results could be obtained.

We think that models with high capacity (CPH-DNNs) readily in themselves cannot improve performance, because they suffer from overfitting. In order to fight overfitting, we idealise that higher models must integrate survival-directed data projection or attention mechanisms and implement robust automatic regularisation procedure through classical means like hyperparameter search or more complex regularisation modules like diet-networks. Missing those important modules, high-capacity models like DNNs working with large dimensionality input like GE data will encounter the same CPH overfitting pitfalls as demonstrated in this paper. Ideal models should allow correct integration of clinical annotations as well, because we have seen that multivariate models tend to perform better if they are available.

An important feature of the CPH is our capacity to interpret the model’s parameters once it is trained, making it an interpretable system. We have chosen not to explore this facet of the algorithm, but we propose to explore this feature further in later work. This should allow the retrieval of the important genetic features of a powerful model and link them to our current understanding of AML. Ideally, to provide useful insight on the biology of AML, survival computer systems from GE data must be robust and interpretable.

## Acknowledgements

The results shown here are in whole or part based upon data generated by the TCGA Research Network: https://www.cancer.gov/tcga.

## References

Bengio, Yoshua, Ian Goodfellow, and Aaron Courville. (2017). Deep learning (MIT press, Vol. 1).

Boutros, P. C., Lau, S. K., Pintilie, M., Liu, N., Shepherd, F. A., Der, S. D., Tsao, M.-S., Penn, L. Z., & Jurisica, I. (2009). Prognostic gene signatures for non-small-cell lung cancer. Proceedings of the National Academy of Sciences of the United States of America, 106(8), 2824–2828. https://doi.org/10.1073/pnas.0809444106

Cox, D. R. (1972). Regression Models and Life-Tables. Journal of the Royal Statistical Society: Series B (Methodological), 34(2), 187–202. https://doi.org/10.1111/j.2517-6161.1972.tb00899.x

Davidson-Pilon, C. (2019). lifelines: Survival analysis in Python. Journal of Open Source Software, 4(40), 1317. https://doi.org/10.21105/joss.01317

Döhner, H., Estey, E., Grimwade, D., Amadori, S., Appelbaum, F. R., Büchner, T., Dombret, H., Ebert, B. L., Fenaux, P., Larson, R. A., Levine, R. L., Lo-Coco, F., Naoe, T., Niederwieser, D., Ossenkoppele, G. J., Sanz, M., Sierra, J., Tallman, M. S., Tien, H.-F., … Bloomfield, C. D. (2017). Diagnosis and management of AML in adults: 2017 ELN recommendations from an international expert panel. Blood, 129(4), 424–447. https://doi.org/10.1182/blood-2016-08-733196

Faraggi, D., & Simon, R. (1995). A neural network model for survival data. Statistics in Medicine, 14(1), 73–82. https://doi.org/10.1002/sim.4780140108

Fradkin, D., & Madigan, D. (2003). Experiments with random projections for machine learning. Proceedings of the Ninth ACM SIGKDD International Conference on Knowledge Discovery and Data Mining, 517–522. https://doi.org/10.1145/956750.956812

Hinton, & Van Der Maaten. (2008). Visualizing Data using t-SNE. Journal of Machine Learning, 9, 2579–2605.

Itti, L., & Koch, C. (2001). Computational modelling of visual attention. Nature Reviews. Neuroscience, 2(3), 194–203. https://doi.org/10.1038/35058500

Kaplan, E. L., & Meier, P. (1958). Nonparametric Estimation from Incomplete Observations. Journal of the American Statistical Association, 53(282), 457–481. https://doi.org/10.1080/01621459.1958.10501452

Katzman, J. L., Shaham, U., Cloninger, A., Bates, J., Jiang, T., & Kluger, Y. (2018). DeepSurv: Personalized treatment recommender system using a Cox proportional hazards deep neural network. BMC Medical Research Methodology, 18(1), 24. https://doi.org/10.1186/s12874-018-0482-1

Kleinbaum, D. G., & Klein, M. (2012a). Introduction to Survival Analysis. In D. G. Kleinbaum & M. Klein (Eds.), Survival Analysis: A Self-Learning Text (pp. 1–54). Springer. https://doi.org/10.1007/978-1-4419-6646-9_1

Kleinbaum, D. G., & Klein, M. (2012b). Kaplan-Meier Survival Curves and the Log-Rank Test. In D. G. Kleinbaum & M. Klein (Eds.), Survival Analysis: A Self-Learning Text (pp. 55–96). Springer. https://doi.org/10.1007/978-1-4419-6646-9_2

Lee, C., Yoon, J., & Schaar, M. van der. (2020). Dynamic-DeepHit: A Deep Learning Approach for Dynamic Survival Analysis With Competing Risks Based on Longitudinal Data. IEEE Transactions on Bio-Medical Engineering, 67(1), 122–133. https://doi.org/10.1109/TBME.2019.2909027

Marquis, M., Beaubois, C., Lavallée, V.-P., Abrahamowicz, M., Danieli, C., Lemieux, S., Ahmad, I., Wei, A., Ting, S. B., Fleming, S., Schwarer, A., Grimwade, D., Grey, W., Hills, R. K., Vyas, P., Russell, N., Sauvageau, G., & Hébert, J. (2018). High expression of HMGA2 independently predicts poor clinical outcomes in acute myeloid leukemia. Blood Cancer Journal, 8(8), 68. https://doi.org/10.1038/s41408-018-0103-6

Ng, S. W. K., Mitchell, A., Kennedy, J. A., Chen, W. C., McLeod, J., Ibrahimova, N., Arruda, A., Popescu, A., Gupta, V., Schimmer, A. D., Schuh, A. C., Yee, K. W., Bullinger, L., Herold, T., Görlich, D., Büchner, T., Hiddemann, W., Berdel, W. E., Wörmann, B., … Wang, J. C. Y. (2016). A 17-gene stemness score for rapid determination of risk in acute leukaemia. Nature, 540(7633), 433–437. https://doi.org/10.1038/nature20598

Papaemmanuil, E., Gerstung, M., Bullinger, L., Gaidzik, V. I., Paschka, P., Roberts, N. D., Potter, N. E., Heuser, M., Thol, F., Bolli, N., Gundem, G., Van Loo, P., Martincorena, I., Ganly, P., Mudie, L., McLaren, S., O’Meara, S., Raine, K., Jones, D. R., … Campbell, P. J. (2016). Genomic Classification and Prognosis in Acute Myeloid Leukemia. The New England Journal of Medicine, 374(23), 2209–2221. https://doi.org/10.1056/NEJMoa1516192

Romero, A., Carrier, P. L., Erraqabi, A., Sylvain, T., Auvolat, A., Dejoie, E., Legault, M.-A., Dubé, M.-P., Hussin, J. G., & Bengio, Y. (2017). Diet Networks: Thin Parameters for Fat Genomics. ArXiv:1611.09340 [Cs, Stat]. http://arxiv.org/abs/1611.09340

Trofimov, A., Cohen, J. P., Bengio, Y., Perreault, C., & Lemieux, S. (2020). Factorized embeddings learns rich and biologically meaningful embedding spaces using factorized tensor decomposition. Bioinformatics (Oxford, England), 36(Suppl_1), i417–i426. https://doi.org/10.1093/bioinformatics/btaa488

Tyner, J. W., Tognon, C. E., Bottomly, D., Wilmot, B., Kurtz, S. E., Savage, S. L., Long, N., Schultz, A. R., Traer, E., Abel, M., Agarwal, A., Blucher, A., Borate, U., Bryant, J., Burke, R., Carlos, A., Carpenter, R., Carroll, J., Chang, B. H., … Druker, B. J. (2018). Functional genomic landscape of acute myeloid leukaemia. Nature, 562(7728) 526–531. https://doi.org/10.1038/s41586-018-0623-z

Wang, F., Jiang, M., Qian, C., Yang, S., Li, C., Zhang, H., Wang, X., & Tang, X. (2017). Residual Attention Network for Image Classification. 3156–3164. https://openaccess.thecvf.com/content_cvpr_2017/html/Wang_Residual_Attention_Network_CVPR_2017_paper.html

Zhao, B., Wu, X., Feng, J., Peng, Q., & Yan, S. (2017). Diversified Visual Attention Networks for Fine-Grained Object Classification. IEEE Transactions on Multimedia, 19(6), 1245–1256. https://doi.org/10.1109/TMM.2017.2648498

